# The FACT complex and cell cycle progression are essential to maintain asymmetric transcription factor partitioning during cell division

**DOI:** 10.1101/075135

**Authors:** Eva Herrero, Sonia Stinus, Eleanor Bellows, Peter H Thorpe

**Affiliations:** The Francis Crick Institute, 1 Midland Road, London. NW1 1AT.; Present address: European Research Institute for the Biology of Ageing, University of Groningen, University Medical Center Groningen, Groningen, The Netherlands; Present address: School of Life Sciences, East Drive, University of Nottingham, Nottingham, NG7 2RD

**Keywords:** Ace2, asymmetry, cell cycle

## Abstract

The polarized partitioning of proteins in cells underlies asymmetric cell division, which is an important driver of development and cellular diversity. Like most cells, the budding yeast *Saccharomyces cerevisiae* divides asymmetrically to give two distinct daughter cells. This asymmetry mimics that seen in metazoans and the key regulatory proteins are conserved from yeast to human. A well-known example of an asymmetric protein is the transcription factor Ace2, which localizes specifically to the daughter nucleus, where it drives a daughter-specific transcriptional network. We performed a reverse genetic screen to look for regulators of asymmetry based on the Ace2 localization phenotype. We screened a collection of essential genes in order to analyze the effect of core cellular processes in asymmetric cell division. This identified a large number of mutations that are known to affect progression through the cell cycle, suggesting that cell cycle delay is sufficient to disrupt Ace2 asymmetry. To test this model we blocked cells from progressing through mitosis and found that prolonged cell cycle arrest is sufficient to disrupt Ace2 asymmetry after release. We also demonstrate that members of the evolutionary conserved FACT chromatin-remodeling complex are required for both asymmetric and cell cycle-regulated localization of Ace2.

## Introduction

Asymmetric cell division is a universal feature of life and provides a key mechanism to create different cell types. It is also particularly important in adult stem cells, where asymmetric cell division maintains a stem cell pool while also generating progenitor cells to repair or replace damaged tissue (Neumuller and Knoblich, 2009). Asymmetric cell division utilizes the polarity axis of the cell to align the mitotic spindle such that the plane of cell division is perpendicular to the axis of cell polarity. In this way, polarized proteins are partitioned differentially into the two daughter cells, potentially altering their fates (Neumuller and Knoblich, 2009). Hence, identifying the mechanisms driving the asymmetric proteins distribution via cell polarity is fundamentally important to understand stem cell function and cell fate determination.

There are a number of mechanisms by which proteins can be polarized ranging from external polarity cues to intrinsic positive and negative feedback that can establish polarity determinants (Johnson et al., 2011; Thompson, 2013). The budding yeast *Saccharomyces cerevisiae* divide asymmetrically during every cell division. The mother cell divides by producing a small protrusion, known as a bud that grows to produce a new daughter cell. The asymmetrical distribution of proteins between the mother and daughter cell leads to a range of divergent phenotypes between these two cells. For example, mother cells progressively age with each cell division, senescing after approximately 30 divisions. In contrast, this replicative ageing process is reset in daughters, which are then themselves able to divide ~30 times as new mothers (Denoth Lippuner et al., 2014). Proteins that are not intrinsically polarized can become so during cell division by selective protein localization to either the mother or daughter cell (Yang et al., 2015). This process is typically driven by the activity of upstream, polarized proteins.

One such protein in *S. cerevisiae* is the transcription factor Ace2. Ace2 regulates genes that are important for daughter cell (bud) specification, especially the separation of the daughter cell from the mother cell and G1 delay (Bidlingmaier et al., 2001; Bourens et al., 2008; Colman-Lerner et al., 2001; Di Talia et al., 2009; Dohrmann et al., 1992; Laabs et al., 2003). Sequential action of kinases and phosphatases generates Ace2 asymmetric localization (see Fig. 1 A). *ACE2* is part of the “CLB2 cluster” of genes that is expressed from early M phase (Spellman et al., 1998). During early mitosis, a nuclear localization signal (NLS) within Ace2 is blocked by mitotic cyclin-dependent kinase (CDK) phosphorylation, which causes Ace2 to accumulate symmetrically in the cytoplasm (Dohrmann et al., 1992). During mitotic exit the Cdc14 phosphatase is released into the cytoplasm. Cdc14 removes CDK phosphorylation from the Ace2 NLS allowing Ace2 nuclear entry (Archambault et al., 2004; Mazanka et al., 2008; Sbia et al., 2008). Ace2 accumulates only weakly in both the nascent mother and daughter nuclei because it is actively exported out of the nucleus, due to a nuclear export signal sequence (NES) (Bourens et al., 2008; Jensen et al., 2000). The RAM network kinase Cbk1 phosphorylates the Ace2 NES blocking Ace2 nuclear export (Brace et al., 2011; Mazanka et al., 2008; Mazanka and Weiss, 2010; Sbia et al., 2008). Although the components of the RAM network localize to the bud neck and daughter cortex during telophase, it is still unclear how the RAM-mediated Ace2 accumulation is restricted to the daughter nucleus (Weiss, 2012).

**Figure 1.**
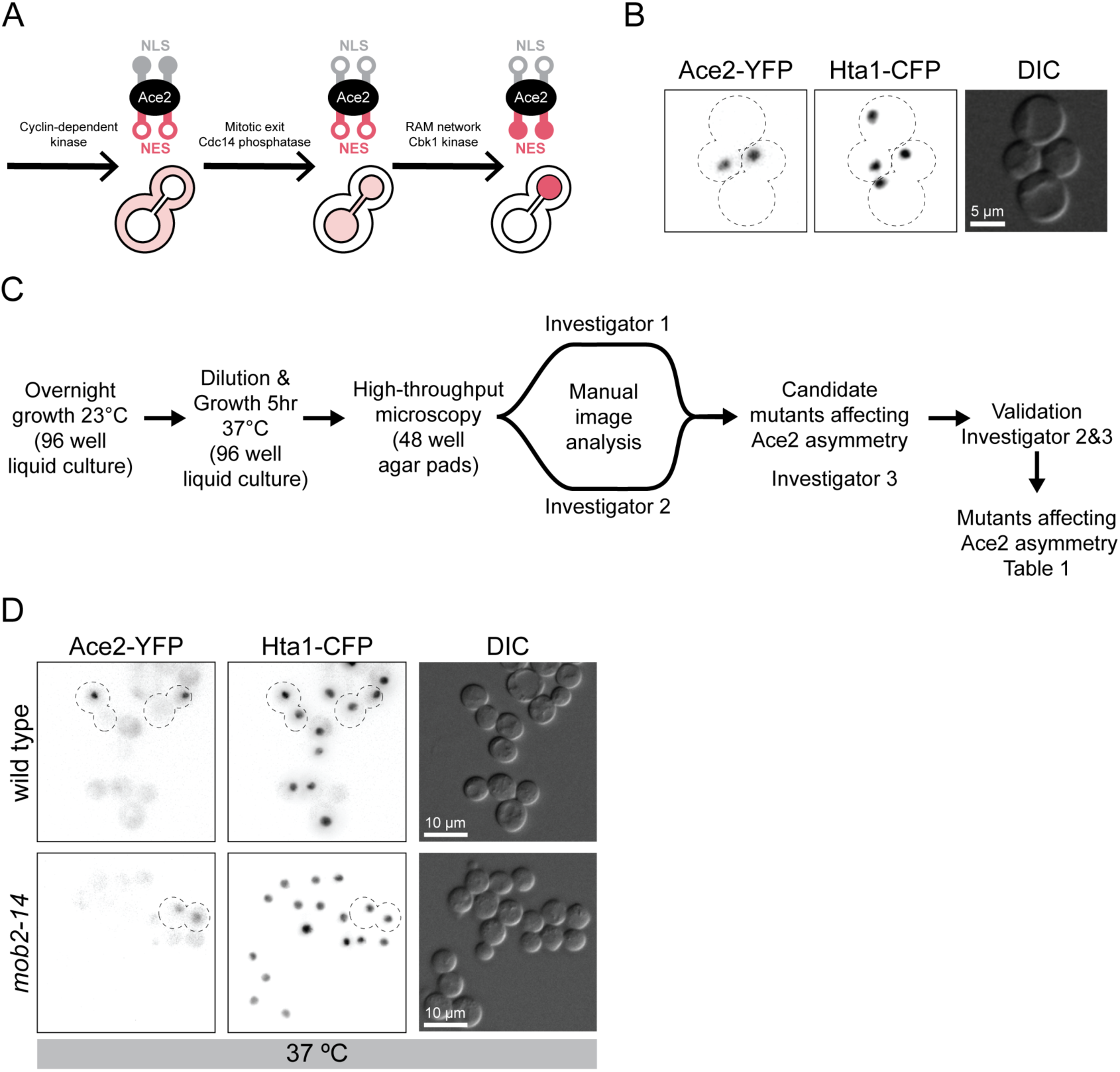
Reverse genetic screen to identify essential genes affecting Ace2 asymmetric localization. (A) Sequential phosphorylation and dephosphorylation controls Ace2 asymmetric localization. (B) Fluorescence imaging of PT31-75D. (C) Fluorescence microscopy screen and image analysis workflow. (D) Fluorescence imaging of wild type and *mob2-14* at the restrictive (37^o^C) temperature. Wild type telophase cells show Ace2-YFP asymmetric localization only in daughter nucleus, whereas *mob2-14* telophase cells show symmetric localization in both mother and daughter nuclei.

The RAM network is one of the yeast Mst/*hippo* or Ndr/LATS signaling systems that are present in most eukaryotic organisms. All members of the RAM network have cell separation defects and loss of asymmetric Ace2 localization (Bidlingmaier et al., 2001; Nelson et al., 2003). Cbk1 is the only “Ndr” kinase in yeast and requires its co-activator Mob2 to function. Cbk1-Mob2 interaction is constitutive throughout the cell cycle. The Cbk1-Mob2 complex also accumulates in the daughter cell nucleus in an Ace2-dependent manner (Colman-Lerner et al., 2001; Weiss et al., 2002). Cbk1 is activated, in part, by the “*hippo”* kinase Kic1, which works with its co-activator Hym1 (Brace et al., 2011; Nelson et al., 2003). However, Kic1 phosphorylation is not sufficient to activate Cbk1. Cdc14 release is also required to remove inhibitory CDK phosphorylation from Cbk1, which in turn allows Cbk1 to interact with Ace2. Hence, Cbk1 phosphorylation of Ace2-NES is restricted to mitotic exit (Brace et al., 2011). However, it is the asymmetric distribution of Cbk1 that is responsible for Ace2’s asymmetry, hence we wanted to ask which proteins regulate this asymmetry. In early G1, Cbk1 keeps Ace2 phosphorylated (Mazanka and Weiss, 2010). Eventually during G1 progression, Ace2 is dephosphorylated and exported into the cytoplasm where a sequestration mechanism involving either Cdk1 or Pho85 prevents Ace2 from re-entering the nucleus (Bourens et al., 2008; Mazanka et al., 2008; Mazanka and Weiss, 2010; Sbia et al., 2008).

A number of cell polarity screens have been performed in yeast. Initially, forward genetic screens were used to identify many of the important genes required for cell polarity, for example *CDC42* (Adams et al., 1990). The creation of arrays of deletions of non-essential genes and overexpression plasmids has enabled the use of reverse genetic approaches to study the cell polarity associated with the mitotic budding process (Styles et al., 2013). For example, the regulation of polarity factors Bni4 and Cdc42 by G1 cyclins was uncovered using synthetic lethality and synthetic dosage lethality screens (Sopko et al., 2007; Zou et al., 2009). Complex haploinsufficiency has been used to identify a role of ESCRT and the proteasome in the regulation of actin filaments and cell polarity (Haarer et al., 2011; Haarer et al., 2007).

The systematic fluorescence imaging of the GFP collection identified hundreds of proteins that localize to sites of polarization such as bud neck, bud tip or actin cytoskeleton (Huh et al., 2003). Moreover, exploring changes in the localization of polarity proteins in a deletion mutant array or gene overexpression array has served to identify regulators of yeast endocytosis (Arlt et al., 2011; Burston et al., 2009). These screens have been highly informative in understanding the regulation of cell polarity and asymmetric cell division. However, systematic loss of function studies with essential genes had not been possible until the creation of collections of hypomorphic and temperature sensitive alleles (Breslow et al., 2008; Li et al., 2011).

Here, we used Ace2 localization as a reporter to test the contribution of essential cellular processes to the maintenance of cell polarity and asymmetry. We performed a fluorescence microscopy screen to assay the localization of Ace2 in an array of temperature sensitive mutants of essential genes. Many of the mutants that affected Ace2 asymmetry are involved in mitotic cell division, and we found that mitotic delay is sufficient to decrease Ace2 asymmetry. In addition, we identified the FACT complex as essential to maintain Ace2 asymmetry, its cell cycle regulated nuclear localization and the localization of the upstream “Ndr” kinase, Cbk1.

## Results

### Identification of essential genes involved in Ace2 asymmetry

To assess which essential genes are required for Ace2 asymmetry, we made use of a collection of temperature-sensitive (ts) mutants in essential genes. This set consists of 787 ts alleles in 497 genes, which covers about half of the essential genes in yeast; each allele is linked to a *KANMX* selectable marker (Li et al., 2011). To assay for Ace2 asymmetry we created a strain (PT31-75D) that encodes Ace2 linked to yellow fluorescent protein (Ace2-YFP) together with a histone H2 peptide (Hta1) fused to cyan fluorescent protein (Hta1-CFP) (Fig. 1 B). Both genes were tagged at the endogenous locus in the genome and remain under the control of their native promoters. This strain shows the characteristic asymmetric distribution of Ace2 in late mitosis (Fig. 1, A and B). Since these two fluorescently marked alleles are linked to selectable markers (nourseothricin resistance for *ACE-YFP* and hygromycin resistance for *HTA1-CFP*), we were able to cross this strain with the ts collection of mutants using synthetic genetic array methodology (SGA) (Tong et al., 2001). This method allowed us to create haploid strains containing the two fluorescent markers in addition to the ts allele, which is linked to a kanamycin resistance gene.. These strains were then assayed for Ace2 asymmetry using fluorescence microscopy after 5 hours of incubation at their restrictive temperature (37°C). Two investigators independently scored the resulting microscope images by visual assessment of whether Ace2 localized asymmetrically and or symmetrically in late mitosis and discrepancies in the resulting scores were resolved by a third investigator (Fig. 1 C). We included a number of wild type controls to ensure that Ace2 asymmetric localization was unaffected by the method (Fig. 1 D). Additionally, we used a ts allele of *MOB2* (*mob2-14*) as a positive control (Fig. 1 D).

Of the 1334 ts strains examined, we were able to score 743 (56%) for Ace2 distribution. There were three main reasons why we could not score the remaining 44%: first, some cells did not grow after SGA, second, some strains did not produce telophase cells after incubation at the restrictive temperature and third, some images were of insufficient quality to score Ace2 distribution. Most of the resulting mutant alleles that were scored as affecting Ace2 asymmetry were then retested individually to remove false positives. Additionally, we retested mutant strains whose genes were functionally or physically associated with the preliminary hits, but we were unable to score in the original screen (33 strains) or they were not a preliminary hit (49 strains). This increased our total number of scored strains to 776 (58% of the total ts alleles, Fig. 2 A). A full list of all the strains and alleles screened plus their Ace2 phenotype is listed in Table S1. We found that 102 strains showed evidence of symmetry breaking at the restrictive temperature (Fig. 2 B, Table S1), which comprise mutations in 81 genes (Fig. 2 C, Table 1). We divided the 81 genes into three phenotypic classes according to the localization of Ace2 (Fig. 2 D). Class I is the largest phenotypic category (90% of mutant genes) and represents a partial phenotype where some cells have wild type Ace2 asymmetric localization while other cells show symmetric Ace2 localization in mother and daughter cell (see Fig. 2 E for example). Two Class II mutants show a complete loss of asymmetric localization of Ace2, and Ace2 was present in all cell nuclei regardless of cell cycle stage (Fig. 2 F). Finally, six Class III mutants show symmetric localization of Ace2 that was restricted to cells in late mitosis (Fig. 2 G). We examined collectively the cellular role of the genes whose mutants affect Ace2 localization (Table 1). A significantly large proportion of them are involved in chromosome segregation (38% of the genes, Fisher’s exact Test p=2×10^−10^, Fig. 2 H). Other represented functions are polarity, chromatin remodeling, transcription and RNA processing, and cell cycle regulation (Fig. 2 H, Table 1).

**Table 1.** List of genes found to affect Ace2-YFP asymmetry Fluorescence microscopy images of *spt16-AID* cells grown for 5 hours in ethanol or auxin (A) Swi5-GFP localization is not altered in Spt16 depleted cells. (B) Cbk1-GFP and (C) Mob2-GFP are mislocalized in Spt16 depleted cells. These images were deconvolved for increased clarity. (D) Ace2-YFP localizes to most nuclei in Spt16 depleted cells both in CBK1 (top panel) and *cbk1*Δ (bottom panel). (E) Experimental setup of G1 depletion of *Spt16-AID*. (F) Fluorescence microscopy of *spt16-AID* cells after G1 release.

| Protein | *ts mutant* | Molecular function |  | Protein | *ts mutant* | Molecular function |  |
| --- | --- | --- | --- | --- | --- | --- | --- |
| Class I |  |  |  | Class I |  |  |  |
| Apc5 | *apc5-CA-Paps* | Cell cycle regulation | * | Cdc7 | *cdc7-4* | DNA Replication | * |
| Cdc27 | *cdc27-2* | Cell cycle regulation | * | Cdc21 | *cdc21-ts* | DNA Replication |  |
| Cdc28 | *cdc28-13* | Cell cycle regulation | * | Psf1 | *psf1-1* | DNA Replication |  |
| Cdc20 | *cdc20-2, cdc20-3* | Cell cycle regulation | * | Kre5 | *kre5-ts2* | Metabolism |  |
| Bet2 | *bet2-1* | Cell transport |  | Krr1 | *krr1-18* | Nucleolar and ribosome |  |
| Sed5 | *sed5-1* | Cell transport |  | Nog2 | *nog2-1* | Nucleolar and ribosome |  |
| Sft1 | *sft1-15* | Cell transport |  | Nop2 | *nop2-5, nop2-9* | Nucleolar and ribosome |  |
| Abf1 | *abf1-102* | Chromatin remodelling |  | Nop7 | *nop7-1* | Nucleolar and ribosome |  |
| Rsc8 | *rsc8-ts16* | Chromatin remodelling |  | Rrp5 | *rrp5-delta6* | Nucleolar and ribosome |  |
| Swd2 | *swd2-1* | Chromatin remodelling |  | Arp3 | *arp3-G302Y* | Polarity |  |
| Tel2 | *tel2-15* | Chromatin remodelling |  | Cdc24 | *cdc24-5* | Polarity | * |
| Ndc10 | *ndc10-1* | Chromosome segregation | * | Cdc43 | *cdc43-2* | Polarity |  |
| Cdc14 | *cdc14-8* | Chromosome segregation | * | Exo70 | *exo70-20/37* | Polarity |  |
| Cdc31 | *cdc31-2* | Chromosome segregation | * | Pan1 | *pan1-4* | Polarity | * |
| Cse4 | *cse4-1* | Chromosome segregation | * | Sec23 | *sec23-1* | Polarity |  |
| Dad2 | *dad2-9* | Chromosome segregation | * | Cdc34 | *cdc34-1* | Protein degradation | * |
| Dam1 | *dam1-1, dam1-19* | Chromosome segregation | * | Met30 | *met30-6* | Protein degradation |  |
| Duo1 | *duo1-2* | Chromosome segregation | * | Pre2 | *pre2-75* | Protein degradation |  |
| Eco1 | *eco1-1* | Chromosome segregation | * | Rpt6 | *rpt6-20* | Protein degradation |  |
| Esp1 | *esp1-1* | Chromosome segregation | * | Gpi13 | *gpi13-5* | Protein modification |  |
| Ipl1 | *ipl1-2* | Chromosome segregation | * | Sec53 | *sec53-6* | Protein modification |  |
| Mif2 | *mif2-3* | Chromosome segregation | * | Cdc39 | *cdc39-1* | Transcription and RNA |  |
| Mps1 | *mps1-1* | Chromosome segregation | * | Hts1 | *hst1-1* | Transcription and RNA |  |
| Mtw1 | *mtw1-ts* | Chromosome segregation | * | Prp19 | *prp19-1* | Transcription and RNA |  |
| Nbp1 | *nbp1-1* | Chromosome segregation |  | Prp2 | *prp2-1* | Transcription and RNA |  |
| Nnf1 | *nnf1-17, nnf1-48, nnf1-77* | Chromosome segregation | * | Rse1 | *rse1-1* | Transcription and RNA |  |
| Nsl1 | *nsl1-5, nsl1-6* | Chromosome segregation | * | Slu7 | *slu7-ts2* | Transcription and RNA |  |
| Nuf2 | *nuf2-61* | Chromosome segregation | * | Ssl2 | *ssl2-ts* | Transcription and RNA |  |
| Pds1 | *pds1-128* | Chromosome segregation | * | Ssu72 | *ssu72-2* | Transcription and RNA |  |
| Sgt1 | *sgt1-3, sgt1-5* | Chromosome segregation |  | Yhc1 | *ych1-1* | Transcription and RNA |  |
| Sli15 | *sli15-3* | Chromosome segregation |  | Class II |  |  |  |
| Smc1 | *scm1-1* | Chromosome segregation | * | Pob3 | *pob3-7, pob3-L78R* | Chromatin remodelling |  |
| Smc3 | *scm3-42* | Chromosome segregation | * | Spt16 | *spt16-1* | Chromatin remodelling |  |
| Spc110 | *spc110-220* | Chromosome segregation | * | Class III |  |  |  |
| Spc24 | *spc24 4-2* | Chromosome segregation | * | Crm1 | *crm1-1* | Nuclear transport |  |
| Spc25 | *spc25-1* | Chromosome segregation | * | Rna1 | *rna1-1* | Nuclear transport |  |
| Spc29 | *spc29-20* | Chromosome segregation |  | Srm1 | *srm1-ts* | Nuclear transport |  |
| Spc34 | *spc34 41-1* | Chromosome segregation | * | Yrb1 | *yrb1-51* | Nuclear transport |  |
| Stu1 | *stu1-12, stu1-5, stu1-6, stu1-7* | Chromosome segregation | * | Hym1 | *hym1-15* | Polarity |  |
| Stu2 | *stu2-11, stu2-13* | Chromosome segregation | * | Mob2 | *mob2-14, mob2-19, mob2-22, mob2-28, mob2-36, mob2-38* | Polarity |  |
| Dbf4 | *dbf4-2, dbf4-3, dbf4-ts* | Chromosome segregation | * |  |  |  |  |
| Smt3 | *smt3-42* | Chromosome segregation | * |  |  |  |  |
| Cdc6 | *cdc6-1* | DNA Replication | * |  |  |  |  |

\* genes in Gene Ontology category "mitotic cell process"

**Figure 2.**
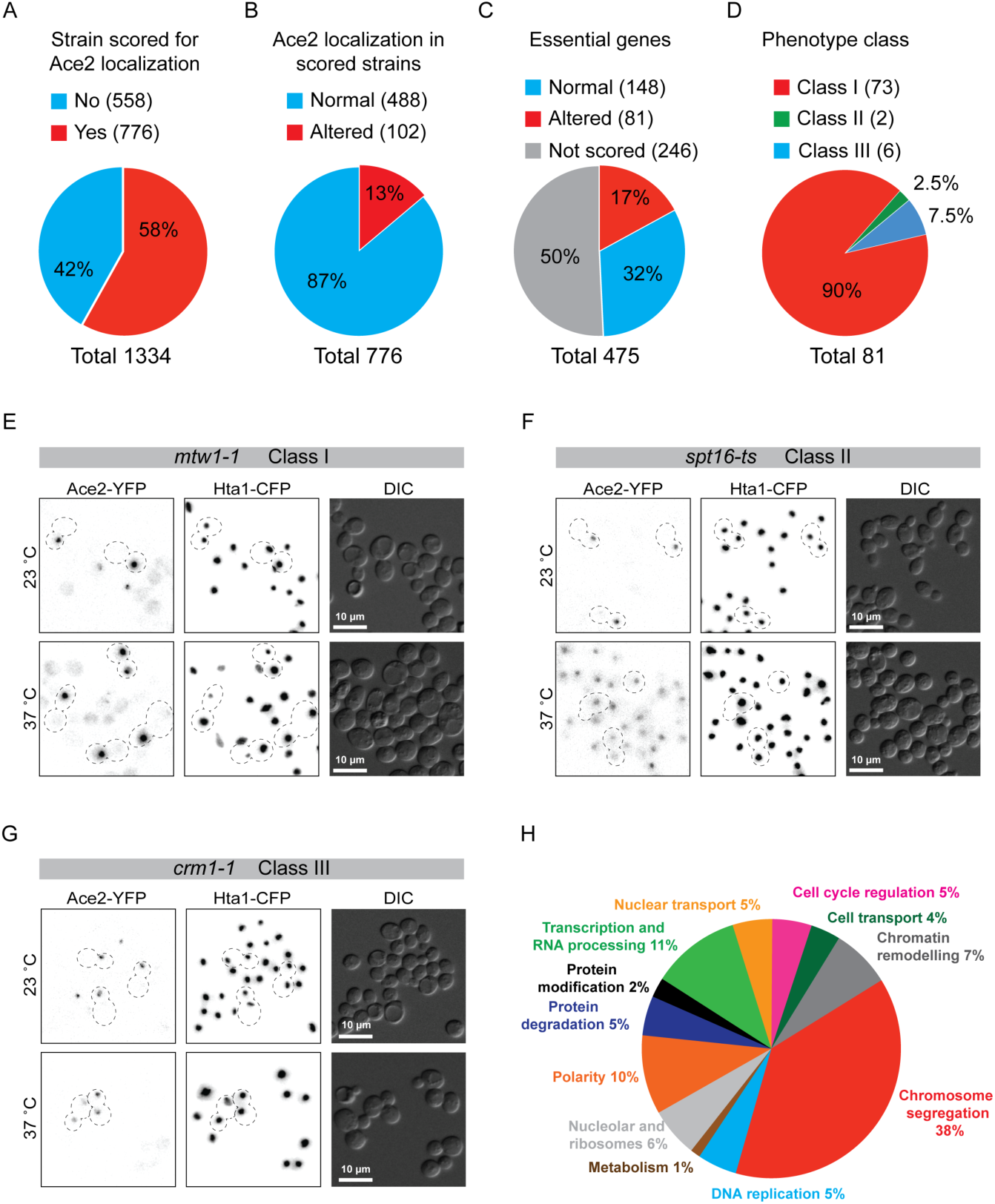
Results from Ace2-YFP reverse genetic screen. (A-D) Summary of screen results: (A) Proportion of total strains scored for Ace2-YFP localization, (B) Characterization of Ace2 localization, (C) Classification of essential genes according to Ace2 localization phenotype, and (D) Phenotypic classification of altered Ace2 localization. (E) Fluorescence microscopy imaging of the Class I mutant *mtw1-1*. At 37 ^o^C both symmetric and asymmetric telophase cells were observed. (F) Fluorescence microscopy imaging of Class II mutant *spt16-ts.* At 37 ^o^C most cells show Ace2-YFP nuclear localization regardless of cell cycle stage. (G) Fluorescence microscopy imaging of Class III mutant *crm1-1*. At 37 ^o^C *crm1-1* telophase cells show symmetric Ace2-YFP localization. (H) Cellular function of hits found to alter Ace2-YFP asymmetric localization. See Table 1.

When validating the Ace2 phenotypes using complementation we noticed that some colonies derived from SGA were ts despite containing a complementing gene for their mutant ts allele. Further investigation showed that the *HTA1-CFP* allele results in a ts phenotype specifically in the W303 genetic background, but not in the BY4741 background. We were able to show that the *SSD1* polymorphism (*ssd1-d*) present in W303 but not in BY4741, which encodes for a truncated Ssd1 protein with no function (Uesono et al., 1997), is responsible for the W303 ts phenotype of *HTA1-CFP* cells (Fig. S1 A). We also tested whether the *SSD1* allele could affect Ace2-YFP asymmetry. We found that *ssd1-d* strains grown at 37 ^o^C have more telophase cells with no Ace2 in either mother or daughter nuclei. However, the percentage of symmetric cells (Ace2-YFP in both mother and daughter nuclei) was similar between *SSD1-V* and *SSD1-D* strains (Fig. S1 B).

### Class III mutants are know regulators of Ace2 asymmetry

As previously reported, *crm1-1* cells showed strong symmetric localization of Ace2 in anaphase-telophase cells (Fig. 2 G) (Bourens et al., 2008; Jensen et al., 2000). We also found mutants of three additional components of the nuclear export machinery - Rna1, Yrb1 and Srm1 - produce a similar phenotype to *crm1-1* (Fig. S2 A and Table 1), as well as alleles encoding two components of the RAM signaling network that regulates Ace2 distribution, Hym1 and Mob2 (6 separate alleles) (Fig. S2 B and Table 1). The other members of this network, Kic1, Tao3 and Cbk1, are not represented in the ts collection. Therefore, Class III mutants validated the ability of our screen to find regulators of Ace2 asymmetry.

### Class I mutants are enriched in mitotic cell cycle processes

#### Mtw1 complementation

One of the Class I mutants is *mtw1-1* (Fig. 2 E). Mtw1 is a structural protein of the kinetochore. To confirm that *mtw1-1* mutation specifically caused this phenotype, we imaged *mtw1-1* mutant cells and *mtw1-1 MTW1* cells (where the mutant allele is complemented with a wild type copy of *MTW1* integrated at the *URA3* locus) at permissive (23 and 30°C) and restrictive (35 °C) temperatures (see Fig. 3 A for 35 °C images), and quantified the loss of asymmetry phenotype. We used Ace2-YFP using fluorescence signal as a surrogate for Ace2 concentration in both mother and daughter nuclei of telophase cells, and assigned cells to three different categories: cells with Ace2 only in one nucleus (asymmetric), cells with Ace2 in both mother and daughter nucleus (symmetric) and cells with no Ace2 in either nuclei (Fig. 3 B). We found that in the *mtw1-1* strain there was a reduction of cells with Ace2 in one nucleus (from 72% to 42%, Fisher’s exact test *p*=0.007), and an increase in the number of cells with Ace2 in both mother and daughter nuclei (from 13% to 50%, Fisher’s exact test *p*=0.002) at the restrictive temperature (35 °C) (Fig. 3 B). The loss of asymmetry phenotype was rescued in *mtw1-1 MTW1* cells, thus confirming that *mtw1-1* mutation specifically caused the phenotype (Fig. 3, A and B). We found that the cumulative fluorescence intensity of mother and daughter nuclear Ace2-YFP (Mother + Daughter nuclear Ace2-YFP) was significantly higher in *mtw1-*1 symmetric cells than in asymmetric cells (Fig 3 C).

**Figure 3.**
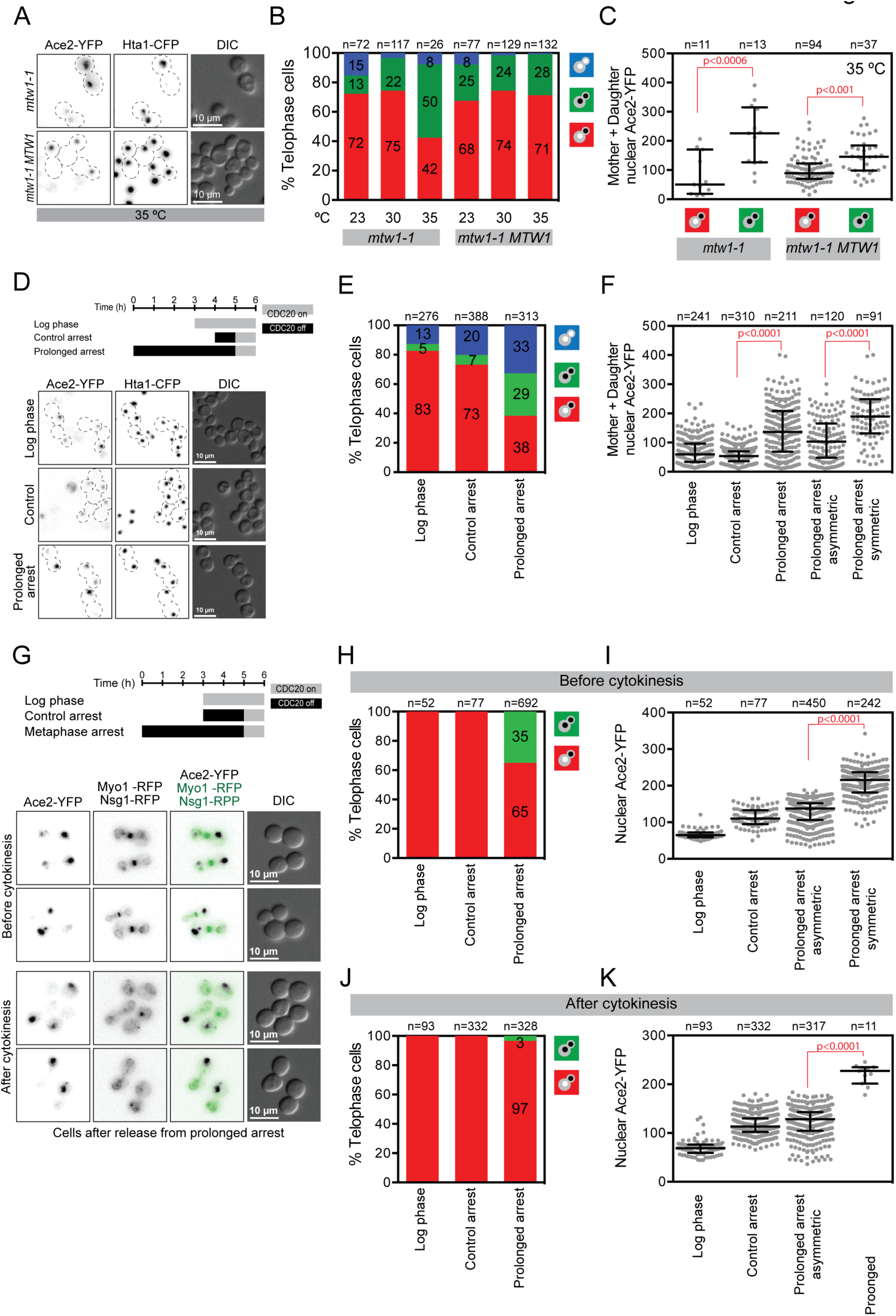
Slow cell cycle progression decreases Ace2-YFP asymmetric localization. (A) Fluorescence microscopy imaging of *mtw1-1* mutant complementation. At 35 ^o^C the partial asymmetry phenotype of *mtw1-1* is rescued by inserting a wild type copy of *MTW1* into the *URA3* locus. (B) Quantitation of Ace2-YFP asymmetry of *mtw1-1* and *mtw1-1 MTW1* cells at three different temperatures. (C) Mother + daughter nuclear Ace2-YFP fluorescence intensity. The plot shows the median and quartiles (error bars). Median (lower quartile 25%, upper quartile 75%): *mtw1-1* asymmetric 50 (19, 171), *mtw1-1* symmetric 226 (126, 315), *mtw1-1 MTW1* asymmetric 89 (70, 123), *mtw1-1 MTW1* symmetric145 (98, 184). (D) Prolonged metaphase arrest experimental set up and fluorescence microscopy imaging. (E) Quantitation of Ace2-YFP asymmetry after prolonged arrest. (F) Mother + daughter nuclear Ace2-YFP fluorescence intensity. The plot shows the median and quartiles (error bars). Median (lower quartile 25%, upper quartile 75%): Log phase 60 (34, 97), Control arrest 54 (36, 70), Prolonged arrest all telophase cells 137 (68, 208), Prolonged arrest asymmetric 103 (48, 165), Prolonged arrest symmetric 190 (132, 249). (G) Prolonged metaphase arrest experimental set up and fluorescence microscopy imaging of cells released from prolonged metaphase arrest before and after cytokinesis. (H) Quantitation of Ace2-YFP asymmetry of cells before cytokinesis from experiment in panel G. (I) Mother + daughter nuclear Ace2-YFP fluorescence intensity of cells before cytokinesis from experiment in panel G. The plot shows the median and quartiles (error bars). Median (lower quartile 25%, upper quartile 75%): Log phase 65 (60, 72), Control arrest 110 (95, 133), Prolonged arrest asymmetric 138 (106, 152), Prolonged arrest symmetric 210 (181, 237). (J) Quantitation of Ace2-YFP asymmetry of cells after cytokinesis from experiment in panel G. (I) Mother + daughter nuclear Ace2-YFP fluorescence intensity of cells after cytokinesis from experiment in panel G. The plot shows the median and quartiles (error bars). Median (lower quartile 25%, upper quartile 75%): Log phase 69 (59, 76), Control arrest 113 (102, 130), Prolonged arrest asymmetric 128 (105, 143), Prolonged arrest symmetric 227 (201, 235). P values in panels C, F, I and K calculated with Kruskal-Wallis Test.

Since 90% of our hits were Class I mutants, we performed a Gene Ontology enrichment analysis within Class I genes (GOrilla) (Eden et al., 2009). We found significant enrichment, 2.6 fold, for genes involved in “mitotic cell cycle process” when comparing the Class I genes to all the genes represented in the ts collection (enrichment *p*-value 3.8×10^−11^). Among these genes, 12 encode kinetochore components such as Mtw1 (see above), spindle and spindle-pole body components such as Spc110 and mitotic regulators such as Cdc27, Ipl1 and Cdc20 (Table 1).

### Metaphase arrest leads to partial loss of asymmetry

A common feature of all of these cell cycle related mutants is that they will likely disrupt the normal progression of the cell cycle. It has been shown that prolonged mitotic delay perturbs asymmetry of acentric DNA (Gehlen et al., 2011) and changes in nuclear shape can decrease the amount of Ace2 asymmetry (Boettcher et al., 2012). Thus we hypothesized that, in the Class I mutants, Ace2 asymmetry is affected by delayed progression through mitosis. To test this hypothesis, we induced a defined mitotic delay and asked whether Ace2 distribution was affected in late anaphase. First, we used depletion of the Cdc20 protein to arrest cells prior to the completion of mitosis. A *CDC20* allele under the control of a *MET1* promoter is transcriptionally repressed by the addition of methionine, which arrests cells in metaphase (Uhlmann et al., 2000). We created a strain that includes the *MET1pr-CDC20* allele together with alleles encoding the tagged versions of Ace2 and Hta1. Cells were arrested in metaphase for either one hour (Control arrest) or five hours (Prolonged arrest) and then released for one hour by transfer into methionine-deficient growth medium, thus allowing cells to enter anaphase and progress to telophase and then imaged them (see Fig. 3 D for example). We quantitatively measured the level of asymmetry of Ace2 as described previously. A short (1 hour) metaphase arrest (control arrest) produced no detectable defect in Ace2 asymmetry when compared with log phase cells (Fig. 3 E). In contrast, we found that prolonged metaphase arrest caused a significant reduction in the proportion of Ace2 asymmetric cells (Fisher’s exact test *p*=9×10^−21^) and an increase in symmetric cells (Fisher’s exact test *p*=7×10^−6^) when compared with the control arrest (Fig. 3 E). Moreover, we found that prolonged metaphase arrest causes increased cumulative levels of Ace2-YFP in telophase cells (Mother + Daughter nuclear Ace2-YFP). Specifically, symmetric cells had higher levels of nuclear Ace2 than asymmetric cells (Fig. 3 F). In contrast, G1 cell cycle delay using alpha-factor did not affect Ace2 asymmetry (Fig. S4). The increase in symmetric cells caused by prolonged metaphase arrest was similar to that in the *mtw1-1* mutant (Figs. 3 B and 3 E). Both *mtw1-1* mutant and prolonged arrest increased the levels of nuclear Ace2-YFP in symmetric cells (Fig. 3, C and F). These data confirm our hypothesis that a delay in mitosis induces a loss of Ace2 asymmetry and it likely explains the Ace2 phenotype found in our primary screen with many of the Class I cell cycle mutants (such as *mtw1, ipl1-1 and spc110-220*).

Ace2 asymmetric localization precedes cytokinesis and persists until G1 (Boettcher et al., 2012; Mazanka and Weiss, 2010). We therefore tested whether there was a specific time when the loss of asymmetry caused by metaphase arrest was manifested. We imaged cells expressing Ace2-YFP, Nsg1-RFP (nuclear envelope) and Myo1-RFP (bud neck) after a control or prolonged metaphase arrest (Fig. 3 G). We quantified symmetric and asymmetric cells before and after cytokinesis, as determined by the disappearance of Myo1-RFP from the bud neck (Fig. 3 G) (Mazanka and Weiss, 2010). Before cytokinesis, there was a significant increase in symmetric cells when subjected to the prolonged arrest (from 0% to 35% asymmetric cells, Fisher’s exact test *p*=3×10^−14^, Fig. 3 H). In contrast, after cytokinesis the majority of cells were asymmetric with Ace2 only in the daughter nucleus (97% asymmetric cells, Fig. 3 J). We again found that symmetric cells had significantly higher levels of nuclear Ace2-YFP than asymmetric cells (Fig. 3, I and K). These data show that Ace2 asymmetry is restored after cytokinesis. Taken together, our data show that prolonged mitosis caused by cell cycle mutations or mitotic arrest decreases Ace2 asymmetry and increases nuclear Ace2-YFP levels.

### The FACT complex is required for Ace2 asymmetry

Only two mutant genes were identified as completely abolishing Ace2 asymmetry in all cells, *SPT16* and *POB3* (Class II). We identified a single allele of *SPT16*, *spt16-1*, and two independent alleles of *POB3*, *pob3-7*, *pob3-L78R*. These two genes encode the two heterodimeric components of the FACT chromatin remodeling complex (Belotserkovskaya et al., 2003; Formosa et al., 2001; Hainer et al., 2012; Ransom et al., 2009; Xin et al., 2009). When we shifted *spt16-1* cells to the restrictive temperature (Fig. 2 F) most cell nuclei contained Ace2 regardless of cell cycle stage (13% at 23^o^C vs. 96% at 37^o^C, Fig. 4 A). We quantified the levels of Ace2 in the nuclei using fluorescence image analysis and found that Ace2 levels are typically less than those of the daughter nuclei at the permissive temperature (Fig. 4 B). To confirm that these proteins are required for Ace2 asymmetry we created a degron allele of *SPT16* (*SPT16-AID*). This construct incorporates a C-terminal addition containing the target site for the AFB2 E3 ubiquitin ligase, whose interaction is dependent upon the presence of auxin (Morawska and Ulrich, 2013; Nishimura et al., 2009). This allele was engineered at the endogenous locus. We found that the total levels of Spt16 are depleted one hour after auxin addition and cells are not viable in the long term (Fig. 4 C). This strain allowed us to test whether we could recapitulate the loss of Ace2 asymmetry seen with the *spt16-1* strain. We incubated the cells with auxin (or ethanol as a control) for 5 hours and assessed the status of Ace2-YFP (Fig. 4 F). Similarly to *spt16-1* at the restrictive temperature most cells grown with auxin contained Ace2 in the nucleus (13% with ethanol vs. 96% with auxin, Fig. 4 D) and at levels that are consistent with those of *spt16-1* (Fig. 4 E). Two of the three *POB3* ts alleles, *pob3-7* and *pob3-L78R* showed a complete loss of asymmetry at the restrictive temperature (Figs. 4 I and S4). However, a third allele *pob3-Q308K* did not show a phenotype (Fig. S4). Further investigations revealed that *pob3-Q308K* was a diploid heterozygous strain containing a wild type *POB3* allele. For *pob3-7*, the proportion of cells with Ace2 in the nucleus was increased from 16% at 23C to 90% at 37C (Fig. 4 G), consistent with the phenotype of *spt16-ts* (Fig. 4 A) and *spt16-AID* (Fig. 4 D). The amount of Ace2-YFP fluorescence in the nucleus was also decreased at the restrictive temperature (Fig. 4 H) although less than in *spt16-ts* (Fig. 4 B), suggesting a less severe phenotype. To confirm *pob3-7* mutation was responsible for the loss of asymmetry phenotype, we complemented the mutant strain with a wild type *POB3* allele cloned into the *URA3* locus. We found that the complemented strain restored wild type Ace2-YFP asymmetric localization and nuclear Ace2 levels at the restrictive temperature (Fig. 4, J-L), thus confirming *pob3-7* mutation as the cause of Ace2 loss of asymmetry.

**Figure 4.**
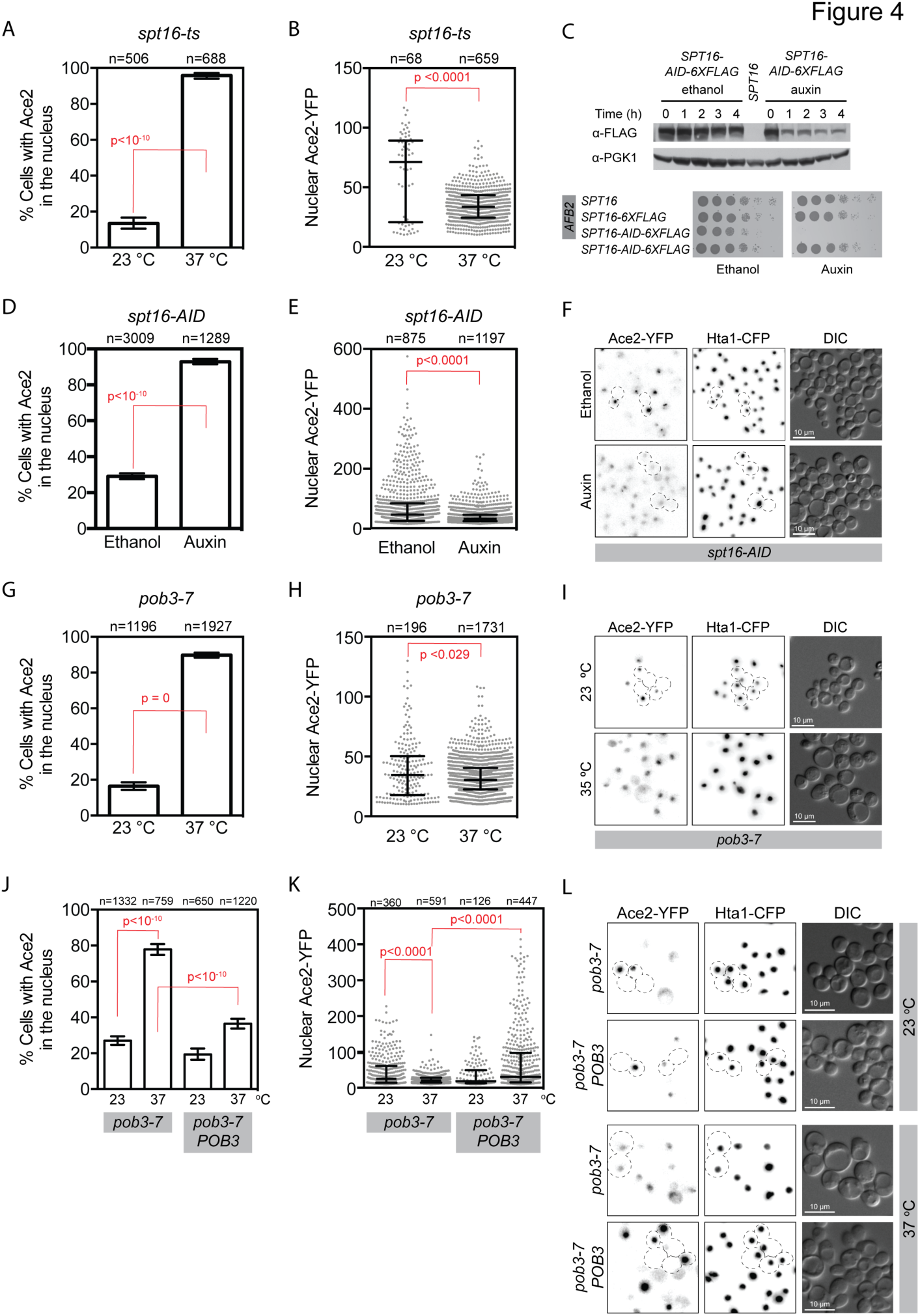
Class II mutants: the two members of the FACT complex Spt16 and Pob3 are required for Ace2 asymmetric localization. (A) Percentage of *spt16-ts* cells with Ace2-YFP in the nucleus at 23 ^o^C and 37 ^o^C. (B) Nuclear Ace2-YFP fluorescence intensity of *spt16-ts* cells. The plot shows the median and quartiles (error bars). Median (lower quartile 25%, upper quartile 75%): 23 ^o^C 71 (21, 89), 37 ^o^C 34 (25, 44). (C) Auxin-dependent degradation of *spt16-AID-6XFLAG* protein, and spot test that shows that *SPT16-AID-6XFLAG AFB2* cells are not viable in the presence of auxin. (D) Percentage of *spt16-AID* cells with Ace2-YFP in the nucleus. (E) Nuclear Ace2-YFP fluorescence intensity of *spt16-AID* cells. The plot shows the median and quartiles (error bars). Median (lower quartile 25%, upper quartile 75%): Ethanol 46 (25, 84), Auxin 31 (24,45). (F) Fluorescence microscopy images of *spt16-AID* cells grown for 5 hours in ethanol or auxin. (G) Percentage of *pob3-7* cells with Ace2-YFP in the nucleus. (H) Nuclear Ace2-YFP fluorescence intensity of *pob3-7* cells. Median (25%, 75% quartiles): 23 ^o^C 35 (18, 50), 37 ^o^C 30 (23, 40). (I) Fluorescence microscopy images of *pob3-7* cells. (J) Percentage of *pob3-7* and *pob3-7 POB3* cells with Ace2-YFP in the nucleus: *pob3-7* 23 ^o^C 27%, *pob3-7* 37 ^o^C 78%, *pob3-7 POB3* 23 ^o^C 19%, *pob3-7 POB3* 37 ^o^C 37%. (K) Nuclear Ace2-YFP fluorescence intensity of *pob3-7* and *pob3-7 POB3* cells. The plot shows the median and quartiles (error bars). Median (lower quartile 25%, upper quartile 75%): *pob3-7* 23 ^o^C 25 (14, 61), *pob3-7* 37 ^o^C 20 (15, 28), *pob3-7 POB3* 23 ^o^C 18 (13, 49), *pob3-7 POB3* 37 ^o^C 30 (15, 97). (L) Fluorescence microscopy images of *pob3-7* mutant complementation. Error bars in panels A, D, G and J correspond to 95% confidence interval. P values calculated with Wilcoxon Rank Sum Test (panels B, E, H) and Kruskal-Wallis Test (panel K).

To allow Ace2 into the nucleus the CDK phosphorylation of the NLS must be reversed by Cdc14 (Archambault et al., 2004; Mazanka et al., 2008; Sbia et al., 2008) (Fig. 1 A). We hypothesized that CDK phosphorylation is perturbed in the FACT complex mutants, such that the NLS is always active (dephosphorylated). To test this we used the location of Swi5 as a surrogate. *SWI5* is a paralog of *ACE2* and encodes a transcription factor whose nuclear localization is also regulated by CDK phosphorylation and Cdc14 dephosphorylation of its NLS signal (Sbia et al., 2008). We reasoned that if CDK phosphorylation of the Ace2 NLS is perturbed, then this would also be true for Swi5. We examined the localization of Swi5-GFP in a strain encoding a marker for the nuclear envelope (Nsg1-RFP) and found that depletion of *Spt16-AID* did not perturb Swi5-GFP nuclear recruitment (Fig. 5 A). These data suggest that the upstream pathway of CDK phosphorylation is not disrupted in the FACT mutants.

**Figure 5.**
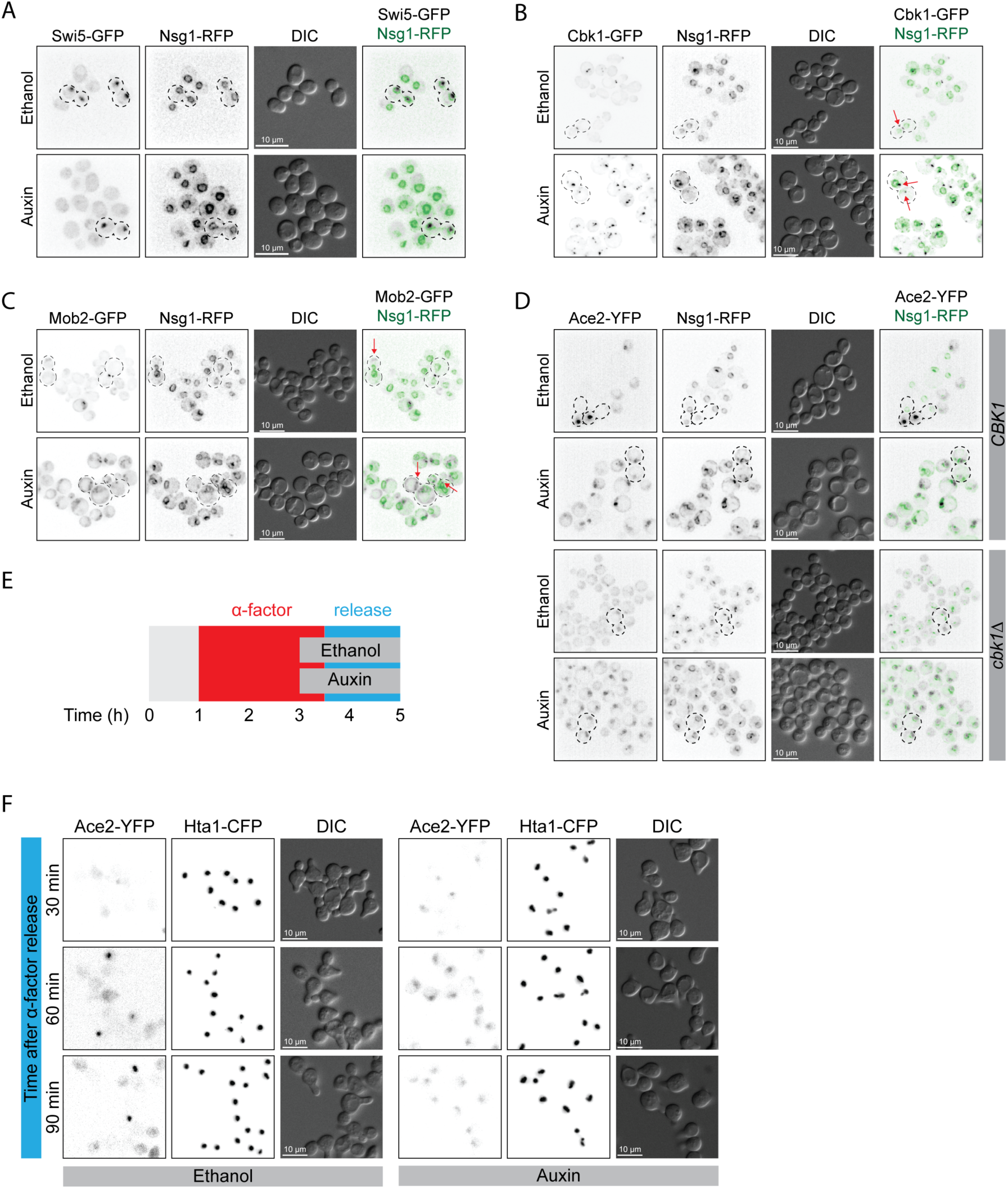
Cbk1-Mob2 complex is mislocalized upon Spt16 depletion.

The next step in Ace2 localization is the phosphorylation of the nuclear export signal by the RAM network, specifically the Cbk1 kinase (Fig. 1 A). We examined the localization of Cbk1-GFP in strains depleted for *Spt16-AID*, also encoding the nuclear envelope marker Nsg1-RFP. Cbk1 is normally restricted to the bud tip in S phase and bud neck during mitosis (Fig. 5 B). However, after depletion of *Spt16-AID* for 5 hours we find that Cbk1 localization is profoundly disrupted with foci of Cbk1 often present outside of the nucleus (Fig. 5 B). Thus a key role for the FACT complex may be to ensure the appropriate distribution of the RAM network. To examine this possibility we looked at the localization of another component of the RAM network, Mob2. Mob2 is required for the specific localization of Cbk1 (Nelson et al., 2003). Like Cbk1, Mob2 is also mislocalized upon depletion of *Spt16-AID* (Fig. 5 C), consistent with the notion that the RAM network localization relies upon the activity of FACT. This suggests that the altered Ace2 localization seen in cells depleted of Spt16 is dependent upon altered Cbk1 localization, which compromises its ability to phosphorylate Ace2. To test this notion we repeated the analysis of Ace2 localization with Spt16 depletion in a *cbk1∆* mutant. In the W303 genetic background *cbk1∆* cells are viable (Bidlingmaier et al., 2001; Nelson et al., 2003), therefore we created Ace2-YFP *cbk1∆* cells with the *SPT16-AID* allele. As previously reported (Mazanka et al., 2008), the *cbk1∆* cells show Ace2 symmetry, but only in telophase cells (Fig. 5 D). Spt16-AID depletion in both wild type and *cbk1∆* cells led to Ace2 present in all nuclei irrespective of the cell cycle stage (Fig. 5 D). These data demonstrate that depletion of the FACT complex produces a more profound Ace2 phenotype in addition to the telophase symmetry seen with *cbk1∆* cells. Thus, the FACT complex may be also interfering with nuclear export of Ace2. To test this notion, we attempted to suppress the *spt16-ts* phenotype by overexpressing nuclear exportin *CRM1 (CRM1-OX*). Crm1 interacts with Ace2-NES and exports Ace2 from the nucleus (Bourens et al., 2008; Jensen et al., 2000). We transformed wild type and *spt16-ts* with a CEN plasmid expressing *CRM1* from the *GAL1* promoter. We grew wild type and *spt16-ts* cells at 37 ^o^C in either raffinose (control) or galactose (*CRM1-OX*) (Fig. S5 A). We found that *CRM1-OX* in wild type and in *spt16-ts* similarly decreased the number of cells with Ace2-YFP in the nucleus, from 51% to 35% and 93% to 83%, respectively (Fig. S5 B). *CRM1-OX* did not change the levels of nuclear Ace2-YFP (Fig. S5 C), these levels were significantly higher in wild type that in *spt16-ts,* as previously shown (Fig. 4 B). These data suggest that the nuclear export of Ace2-YFP is not affected in *spt16-ts*. FACT mutants are inviable and *spt16-ts* cells arrest in G1 at the restrictive temperature (Prendergast et al., 1990), suggesting an essential role of the FACT complex in G1. Ace2 is inactivated during G1 progression when Cbk1 phosphorylation decreases, leading to the export of Ace2 from the nucleus and its sequestration within the cytoplasm (Mazanka and Weiss, 2010). To test whether Spt16 plays a role in Ace2 inactivation during G1, we arrested *SPT16-AID* cells in G1 with α-factor and induced *spt16-AID* depletion by adding auxin before releasing the G1 arrest (Fig. 5 E). Upon G1 release, control cells progressed through the cell cycle and daughter-specific Ace2-YFP localization in telophase cells was visible after 60 minutes (Fig. 5 F Ethanol). In contrast, G1 arrest persisted in *spt16-AID* depleted cells, as previously described (Morillo-Huesca et al., 2010; Prendergast et al., 1990). Interestingly, Ace2-YFP was present in the nuclei of all the cells, suggesting that Ace2 cytoplasmic sequestration is impaired in the absence of Spt16 (Fig. 5 F Auxin).

Taken together, our data demonstrate that the FACT complex is essential to maintain Ace2 asymmetric localization and cell cycle regulation. Although Cdk1 phosphorylation is not altered in Spt16 depleted cells, RAM network Cbk1-Mob2 localization and G1 Ace2 cytoplasmic sequestration are altered in the absence of Spt16.

## Discussion

We have systematically screened a collection of ts mutants of essential genes for those that perturb the asymmetry of a canonical marker of asymmetric cell fate determination, Ace2 (Fig. 1). We found 81 genes whose disruption abrogates Ace2 asymmetric distribution and we grouped these into three phenotypic classes. The vast majority are Class I, which show some but not all telophase cells with Ace2 mislocalized (Fig. 2). This class includes many genes that are expected to disrupt the cell cycle and especially those involved in mitosis such as *MTW1*, *IPL1* and *CDC20*. Other studies have shown that Ace2 asymmetry is perturbed in cells with altered nuclear morphology (Boettcher et al., 2012) and that asymmetric distribution of acentric DNA is altered by delayed mitosis (Gehlen et al., 2011). In line with these studies, we found that a prolonged mitotic arrest is sufficient to abrogate Ace2 asymmetry (Fig. 3). These data together with that described above suggest that prior to the separation of the nuclei in dividing cells, Ace2 can diffuse between the two nascent nuclei, and that a rapid mitosis contributes to the generation of Ace2 asymmetry. We also found that Ace2 asymmetry was restored after cytokinesis, implying that the underlying mechanism (CDK phosphorylation and the RAM network) is functioning normally in these cells. We suggest that Ace2 present in the mother nuclei is exported due to the lack of Cbk1 phosphorylation that only occurs in the daughter (Mazanka et al., 2008). This suggests a mechanism that actively creates the asymmetric distribution of Ace2 (phosphorylation dependent inactivation of NLS and NES signals) by balancing the diffusion of Ace2 along a concentration gradient. Interestingly, prolonged metaphase arrest increased the amount of nuclear Ace2, especially in symmetric cells, and also in Class II mutant *mtw1-1* (Fig. 3). This increase in Ace2 protein is presumably due to an increase of *ACE2* transcription, since *ACE2* expression is restricted to M phase (Spellman et al., 1998). Hence, it is possible that increasing the total amount of Ace2 protein contributes to the perturbation of its asymmetric partitioning.

The Class II mutants were limited to two genes, *SPT16* and *POB3*, which encode the two heterodimeric components of the FACT complex. The FACT complex plays an essential role in chromatin disassembly and reassembly during transcription elongation, and hence contributes to general chromatin structure maintenance (Formosa, 2008; Reinberg and Sims, 2006). In our study, *spt16* and *pob3* mutants completely abolished Ace2 asymmetry and, furthermore, contained Ace2 within the nucleus of cells outside of mitosis (Fig. 4). This phenotype suggests that both CDK and Cbk1 phosphorylation of Ace2 are affected in the Class II mutants causing nuclear accumulation of Ace2 regardless of cell cycle stage and loss of asymmetry, respectively. The downregulation of the G1 cyclins (*CLN1, CLN2* and *CLN3*) in *spt16* mutants leads to a low CDK activity and defects in progression to START, the G1/S transition (Prendergast et al., 1990; Rowley et al., 1991). In the absence of FACT activity, excessive free-histones accumulation leads to the downregulation of *CLN3* (Morillo-Huesca et al., 2010). Cln3 activation of Cdc28 (CDK) is required to activate *CLN1* and *CLN2* expression (Costanzo et al., 2004; de Bruin et al., 2004; Takahata et al., 2009; Wang et al., 2009). Low CDK activity in *spt16* mutant cells could explain the loss of Ace2 cell cycle regulation since Cdc28 phosphorylation of Ace2-NLS prevents Ace2 nuclear import (Archambault et al., 2004; Dohrmann et al., 1992; Mazanka et al., 2008; Sbia et al., 2008). However, we found that Swi5 cell cycle regulation was not altered in *spt16* depleted cells (Fig. 5 A), suggesting that NLS phosphorylation was not primarily affected. Interestingly, we found Spt16 depletion in G1 arrested cells causes nuclear accumulation of Ace2 in all cells (Fig. 5 F), suggesting that Spt16 is required for Ace2 cytoplasmic sequestration. Both CDKs Pho85 and Cdc28 act redundantly to sequester Ace2 in the cytoplasm of daughter cells (Bourens et al., 2008; Mazanka et al., 2008; Mazanka and Weiss, 2010; Sbia et al., 2008). This suggests that elevated levels of nuclear Ace2 seen in Spt16-depleted G1 cells, may be the result of reduced CDK activity in G1, however, our data with Swi5 suggest that CDK activity is not affected in mitosis (Fig 5 A). Moreover, the presence of Ace2 is all cells, both mother and daughters, suggests that Spt16 has additional roles in Ace2 cytoplasmic sequestration. We also found that both RAM network components Cbk1 and Mob2 are mislocalized in an *spt16* mutant (Fig. 5, B and C), suggesting that the loss of Ace2 asymmetry is due to defective Cbk1 phosphorylation of Ace2. Consistently, deletion of *CBK1* (*cbk1*Δ) did not change Ace2 localization phenotype of *spt16* mutant cells (Fig. 5 D).

Taken together, we found that essential processes affect the asymmetric distribution of Ace2 protein. A mitotic delay reduced Ace2 asymmetry but did not compromise Ace2 cell cycle regulation. Diffusion of proteins from one nascent daughter cell to the other is likely prevented by rapid mitosis and cytokinesis. Based on previous studies (Boettcher et al., 2012; Gehlen et al., 2011), we suggest that mitotic delay allows diffusion to break the asymmetry of many different asymmetrically distributed molecules. We found a novel critical role of the FACT complex to maintain the correct localization both Ace2 and RAM network. The FACT complex also affects the nuclear levels of Ace2 in G1 cells. It will be relevant to determine the mechanism by which chromatin remodeling is involved in the localization of the RAM network proteins. Furthermore, it will of interest to investigate whether chromatin-remodeling factors, such as the FACT, play a role in the localization of conserved “*hippo/ndr*” kinases in other eukaryotes.

## Methods

### Strains

A full list of strains is included in Table S1 and a full list of ts alleles tested in the screen is included in Table S2. Yeast medium and growth was performed using standard methods (Sherman). We constructed a strain (PT31-75D) that includes *HTA1-CFP::HYG* and *ACE2-YFP::NAT* in addition to the haploid specific marker *lyp1∆::STE3pr-LEU2*. We crossed this *MATα* strain with the 787 members of the *MAT***a** temperature-sensitive collection *GENEX-TS::KANMX* (Li et al., 2011) using the SGA technology (Tong et al., 2001) and used the ROTOR pinning robot (Singer Instruments, UK) to copy the cells on the different selection media. Diploids were selected on YPD with geneticin (G418) and nourseothricin (NAT), and then sporulated in sporulation media (SPO-SGA). The resulting spores were selected on synthetic medium lacking leucine and supplemented with 50µg/ml thialysine for *MATα* haploids. This media was supplemented with G418, NAT and hygromycin (HYG) to select for fluorescently-tagged and ts alleles.

To generate *SPT16-AID::HYG ura3-1::ADH1pAFB2*, the plasmid pHT453 digested with *StuI* was transformed into the strains containing the appropriate fluorescent reporters. Then, the *AID-6XFLAG::HYG* cassette was amplified from the plasmid pX58 (see primers Table S3). Transformed strains were confirmed by PCR and sequencing. Cell viability was assayed by spotting cells in 500 μM auxin.

### Microscopy

For the screen, cells were prepared for imaging by growth in ~250 µl of liquid synthetic media supplemented with adenine (100 mg/L, +Ade) in 96-well plates overnight at 23˚C. These cultures were then diluted (1:10) and grown at 37˚C for 5 hours. The yeast were imaged on agar pads (Werner et al., 2009) using a 63x, 1.4NA oil immersion objective lens (Carl Zeiss A.G., Germany). Fluorophores were excited with LED-based illumination at 445nm for CFP and 505nm for YFP using appropriate filter sets (47HE for CFP and 46HE for YFP, Carl Zeiss A.G., Germany). Fluorescence images were acquired on a charge-coupled device (CCD) camera (Orca ERII, Hamamatsu Photonics K.K., Japan) with exposure times of 10msec for CFP and 150msec for YFP, the CCD pixels were binned 2x2 for an improved signal to noise ratio.

For other microscopy, cells were grown overnight in the appropriate synthetic media +Ade. On the day of the experiment, cells were diluted to OD 0.3. For experiments using ts strains, cells were grown for 5 hours at permissive or restrictive temperature. For auxin-dependent Spt16-AID depletion experiments, log phase cultures at OD_600_ =0.3 were grown for one hour before adding 500 µM auxin (from 100 mM stock in 100 % ethanol) or only ethanol. Then, cells were grown for 5 hours before imaging. For *CRM1-OX* experiments, cells were grown in synthetic media with 2% raffinose and 0.1 % glucose. An additional 2% galactose was added to induce *CRM1-OX* from the GAL1 promoter. For metaphase arrest experiments, *MET3pCDC20* strains were grown overnight in synthetic media lacking methionine (-Met). The day of the experiment, cultures were diluted to OD 0.3 and grown for one hour. Then, cultures were spun down and resuspended in synthetic media with 2mM methionine (+Met), and grown for the indicated time. To release from the arrest, cultures were washed twice with distilled water and resuspended in –Met. Then cells were grown and imaged every 30 minutes to follow mitotic progression. For imaging cells were mounted on microscope slides with 0.7% LMP agarose in the appropriate synthetic media +Ade, and covered with 0.17 mm glass coverslips. Cells were imaged with a 63x, 1.4NA oil immersion objective lens (Carl Zeiss A.G., Germany). YFP and CFP fluorophores were excited as explained above. Additionally, GFP and RFP were excited with LED-based illumination at 470 nm for GFP and 590 nm for RFP using appropriate filter sets (38HE for CFP and 60HE for RFP, Carl Zeiss A.G., Germany). Images were captured either with a CCD camera (see above) or a CMOS camera (Flash 4.0 lte, Hamamatsu Photonics K.K., Japan).

### Image analysis

For the screen, images were scored manually using Volocity imaging software (Perkin Elmer Inc., USA). Two investigators scored the microscope images independently by visual assessment according to the criteria listed in Table S2, principally to determine whether Ace2 was localized asymmetrically or symmetrically in late mitosis (Fig. 1 D). A third investigator resolved discrepancies in the resulting scores independently to produce a preliminary list of hits. We then retested these preliminary hits and other mutant strains, whose genes were functionally or physically associated with the preliminary hits, but were not scored or did not register as a hit in the original screen.

For quantitation of Ace2-YFP, we used a custom-made protocol in Volocity imaging software. Hta1-CFP fluorescence intensity was used to automatically segment yeast nuclei. A background region was selected by subtracting a 2X dilation from a 4X dilation of the yeast nuclei. For every object (nuclei and background), both CFP and YFP mean fluorescence was measured. Background mean intensity was subtracted from nuclei mean intensity to calculate the corrected nuclear intensity. Telophase cells were manually selected from the images. The corrected Ace2-YFP nuclear intensity of S phase cells was used as a threshold to define nuclei without Ace2-YFP. The presence or absence of Ace2-YFP in either one or both nascent mother or daughter nuclei was used to assign telophase cells to the categories indicated in figures 2 and 3. Cumulative fluorescence intensity between mother and daughter cells of nuclear Ace2-YFP was calculated by the sum of the corrected Ace2-YFP of both nascent nuclei of telophase cells. For the experiment in Figure 3, H-K, automatic nuclei segmentation was achieved by using Ace2-YFP intensity, thus cells without Ace2-YFP either in mother or daughter nuclei are not included in the analysis. Bud-neck localized Myo1-RFP was automatically segmented using RFP fluorescent intensity. The images shown in Figure 5, A-C were deconvolved for increased clarity. We used the deconvolution algorithm in Volocity software (Perkin Elmer, USA), using a simulated point spread function.

### Western blot analysis

Whole cell extracts were prepared as previously described (Olafsson and Thorpe, 2015). 10 μl of protein extract were loaded in a 7.5% acrylamide gel (Biorad). Proteins were transferred into a 0.45 μm supported nitrocellulose membrane (Biorad). Membrane blocking and antibody incubation were performed using Western Blocking reagent (Roche). Anti-FLAG antibody (Sigma, F7425) and anti-GFP antibody (Roche, 11814460001) were used at 1:1000 dilutions. Anti-Pgk1 antibody (Invitrogen, 459250) was used at 1:10000. HRP conjugated anti-Rabbit IgG (Sigma, A0545) and anti-mouse IgG (Abcam, ab97265) were used at 1:100000 and 1:30000 dilution, respectively. Membranes were incubated for 1 min with Lumi-Light Western Blotting substrate (Roche) before film exposure.

## Acknowledgements

We would like to thank Helle Ulrich for the AIS system plasmids and Lisa Berry for critical comments on this manuscript. This work was supported by the Francis Crick Institute which receives its core funding from Cancer Research UK (FC001183), the UK Medical Research Council (FC001183), and the Wellcome Trust (FC001183); by the UK Medical Research Council (MC_UP_A252_1027) and Sonia Stinus was supported by an award from European Union (Unión Europea, Leonardo da Vinci programme), executed by Confebask.

